# Harnessing variation in *Drosophila melanogaster* reveals different routes to increased infection susceptibility over age and sex-dependent association with lifespan

**DOI:** 10.1101/2025.07.07.663438

**Authors:** Barbara Monteiro-Black, Mary-Kate Corbally, Julia Aleksandrowicz, Adam M Taggart, Francesco Langella, Rebecca L Belmonte, Paulina Mika, David F Duneau, Jennifer C Regan

**Affiliations:** Institute of Immunology and Infection Research, University of Edinburgh, Edinburgh, United Kingdom; Center for Ecology, Evolution and Environmental Changes (cE3c) & Global Change and Sustainability Institute (CHANGE), Faculty of Sciences, University of Lisbon (FCUL), 1749-016 Lisbon, Portugal; Department of Pathology, University of Cambridge, Cambridge, England, UK; University of Oxford, Oxford, England, GB; Centre for Reproductive Health, Institute for Regeneration and Repair, University of Edinburgh, United Kingdom

## Abstract

It is widely accepted that susceptibility to infection increases with age, usually attributed to immune system dysregulation. However, factors beyond decreased resistance mechanisms may increase susceptibility, such as changes to physiological or metabolic processes determining overall tolerance, and this may differ between individuals. We hypothesised that age-related increases in susceptibility reflect changes in both immune system function and disease tolerance, with their relative importance varying by population and sex. We assessed age-related susceptibility to bacterial pathogens in an outbred *D. melanogaster* population and 22 isolines. We leveraged this variation to investigate age-related infection susceptibility and lifespan, and to estimate frequencies of declines in resistance or tolerance mechanisms. Susceptibility increased early in life, and was widespread across infections and populations. Despite strong correlation between the sexes in baseline susceptibility and its decrease over age, susceptibility at old age did not correlate. Increased susceptibility was associated with impaired resistance (*bona fide* immunosenescence), but more frequently with reduced tolerance, which could be detected early in life, and remained low. We find susceptibility at old age predicts lifespan in males only. In conclusion, increased infection susceptibility is an early-arising ageing phenotype occurring in both sexes and driving male mortality. Infection susceptibility followed a power-law (Weibull) rather than the exponential (Gompertz) pattern typical of mortality, with each doubling of age approximately doubling susceptibility. We argue that the term “immunosenescence” should be limited to dysregulation of resistance mechanisms, and consideration of population variation, sex, and ageing of non-immune functions is critical to fully understand age-related loss of pathogen control.

## Background

Susceptibility to infection increases with age and this is observed across taxa – in diverse laboratory animal models, wild animal populations, and in humans [1–4]. However, we do not understand what underpins the increase in susceptibility to pathogens over ageing, and despite established sexual dimorphism and population diversity in ageing phenotypes [5–8], we know little about variation in increased infection susceptibility, including sex differences [9–13]. A long-standing assumption is that increased infection risk is underpinned by deterioration of immune functions that act to reduce pathogen burden. However, many functions other than immunity decline with age, potentially before the decline of the immune system itself. The decline of key metabolic, digestive or neuronal functions could potentially result in increased susceptibility to infection [14–16] if they decrease the ability to tolerate the consequences of infections (e.g., increased sensitivity to tissue damage or decreased repair). If mechanisms that influence the ability to tolerate infections suffer a functional decline over age, this challenges resistance-focussed assumptions around age-related increased susceptibility to infections, and would require different therapeutic targets. The scarcity of models that allow for longitudinal experimental analysis means we do not know, in any system, the relative contributions of immune system failure, or reduction in other physiological functions, to increased infection susceptibility at old age [12]. Sexual dimorphisms in immunity, metabolism, and endocrine regulation are likely to shape both resistance and tolerance in distinct ways across the lifespan, including the balance of their decline in older individuals. We set out to test these questions, predicting that a reduced ability to tolerate the consequences of infection will be a major contributor to age-related infection risk, alongside the well-recognised decline in immune tissue function, and that the sexes will show distinct patterns of increased infection susceptibility and correlation with mortality [17].

Host susceptibility is determined by two largely distinct, but non-mutually exclusive, strategies: resistance and disease tolerance [18–21]. Resistance describes the host’s ability to directly limit the pathogen burden, and is determined by the action of immune effectors such as antibodies, antimicrobial peptides (AMPs), reactive oxygen species (ROS) and phagocytic cells. Disease tolerance (hereafter referred to as tolerance) does not directly affect pathogen proliferation, but describes the host’s ability to limit disease severity and maintain health in spite of a dynamic pathogen load [19, 22–24]. Tolerance is a broad term encompassing processes such as the sensitivity to, and repair of, tissue damage [25–27], and maintenance of cellular [28, 29] and metabolic homeostasis [15, 30–32]. Consequently, an increase in susceptibility due to a decrease in tolerance can arise from defects in various physiological processes (i.e., from anything that is not a *bona fide* immune function). Because resistance and tolerance depend on many tissue-level functions and mechanisms, it is impractical to measure them all, and challenging to decide which functions to quantify in order to attribute susceptibility to either strategy. Simply measuring the quantity of investment in a given mechanism may not be informative. For instance, immune cells present in similar numbers may have differing functional efficiencies across individuals, or within an individual over ageing [33]. The efficiency of a mechanism may be a more suitable measure than its quantity, and not necessarily correlated. Immune efficiency depends on a balanced response, the threshold of which is unknown and could be detrimental if exceeded [34–38]. Immune dysfunction over ageing is often linked to reduced regulation leading to over-production, rather than under-production, of immune components, like AMPs [39]. Furthermore, immune mechanisms are operating within a complex system, and a decrease in efficiency of one arm of defence could be compensated by another [40]. Studying the relative contribution of resistance and tolerance to infection susceptibility is further complicated by their connectedness [41, 42]. For example, during an intestinal infection, reduced tissue repair capacity could increase infection-induced damage to the gut barrier. This damage could result in reduced nutrient absorption, impacting the host’s ability to mount an energetically costly immune response, thereby reducing resistance. Work integrating theoretical modelling and empirical testing has suggested that studying pathogen loads at key points of an infection can establish the relative contributions of resistance and tolerance without the need to target specific mechanisms [42–46].

Analysing pathogen load is an ideal way to assess host performance without exhaustively defining parameters: it is the result of pathogen proliferation within the host, which necessarily depends on both the host’s ability to control the proliferation (resistance) and to sustain its impact (tolerance). However, it is extremely challenging to measure in mammalian models, as most infections are within tissues and consequently not easily accessible, and multiple sampling over time is limited by ethical concerns. In addition, the number of hosts per experiment is usually small [47, 48], precluding analysis of several populations, different genotypes, and even of both sexes. Following pathogen load over ageing would be the ideal way to establish the connection between susceptibility and defence strategies – i.e., resistance and tolerance – if these technical challenges could be overcome.

Using insects to address outstanding fundamental questions in immune system ageing is one possible approach [12, 48]. In mammals, age-related increased susceptibility can be attributed to age-related alterations in both the adaptive [1, 49] and innate arms of immunity [50, 51]. Increased susceptibility to infection is also observed in invertebrate systems that possess only innate immunity [52–54]*. D. melanogaster* is a cornerstone genetic model system that has classically been used in the biology of ageing, biogerontology, and immunity fields due to high genetic conservation in relevant signalling pathways [12]. In ageing *D. melanogaster*, the transcription of AMPs, induced by NF-kB signalling pathways, becomes dysregulated [55–57], reflective of inflammatory transcriptional profiles in ageing vertebrates [58, 59]. AMP induction also becomes less specific to bacterial infections in older individuals [60], and macrophage-like hemocytes decline in their phagocytic capacity [61, 62]. In addition, the intestinal epithelium in older individuals becomes disorganised and leaky, harbouring a higher load of potentially pathogenic bacteria [63, 64], a feature of ageing which shows a particularly striking sex dimorphism [65]. Overall, there is a consensus that older flies are more susceptible to infections, including by bacteria, viruses, and fungi [66], suggesting that immune ageing in insects follows a broadly similar pattern to mammals [67], including stark differences between the sexes [68].

In this study, we assessed pathogen load to elucidate which immune strategies (i.e., resistance or tolerance) underpin increased infection susceptibility over age, and whether immune ageing correlates with longevity. We first show that aged individuals from a genetically heterogeneous population have a reduced ability to survive a range of bacterial infections, including Gram-negative and Gram-positive infections through systemic and oral routes, controlled by different immune responses. We next established multiple, large populations from a subset of isogenic lines of the *D. melanogaster* Genetic Reference Panel (DGRP) [69] and challenged them at young and old ages with the natural *D. melanogaster* pathogen *Providencia rettgeri*. We also measured the lifespan of unchallenged flies from a subset of the populations. Population variation, arising from both intrinsic (e.g., genotype, sex) and extrinsic (i.e., environmental) factors, allowed us to measure co-variation in early- and late-life susceptibility to systemic infection, and in lifespan. Increased susceptibility was widespread across populations and sexes, but varied strongly in magnitude. The change in the <u>P</u>athogen <u>L</u>oad <u>U</u>pon <u>D</u>eath (PLUD) with age across the panel enabled us to estimate the relative contribution of immune strategies to increased susceptibility [42]. This showed that, most commonly, increased susceptibility was driven primarily by lowered tolerance. However, in several populations, the increase in susceptibility to systemic infection was mainly due to a loss of resistance – i.e., a decreased control of pathogen load. Tracking PLUD across the life course in a tolerance-deficient population, our findings suggested that, at old age, the driver for increased susceptibility can remain consistent – i.e., tolerance decreased at middle age, and remained low into old age. We showed that there was no association between early-life susceptibility and lifespan in either sex. However, there was a significant negative correlation between late-life susceptibility and lifespan in males, regardless of the existence or direction of sex bias in longevity. Overall, we showed that pathogen load associated with mortality risk can show variable routes to increased susceptibility – with loss of tolerance being a common driver – although death can also be driven by *bona fide* loss of immune function. These findings establish that further work across systems should not only focus on the dysregulation of immune tissue resistance mechanisms, but also on the dysregulation of non-immune functions, and how this loss of tolerance impacts the ability to control infections.

## Results

### Increase in infection susceptibility occurs across range of pathogenic contexts

Susceptibility to bacterial infection is expected to increase over age [70–72]. To investigate its dynamics and define what was a “young” and an “old” individual, we parameterised infections by infecting a single isoline genotype (RAL-821) at weekly intervals through its lifespan (up to 40 days) using the *D. melanogaster* bacterial pathogen, *P. rettgeri* [73–75]. As expected, infection susceptibility increased over ageing relative to 7-day-old individuals for both males and females (Figure 1). Increased infection susceptibility was detected at day 14 in females, where risk trended upwards continually until our final measurement at day 40. In males, risk increased significantly between day 14 and 21, and also trended upwards until 40 days (Figure 1A and A’). To describe the mathematical pattern underlying the dynamics of age-related increase in infection susceptibility [76], we compared five theoretical models of biological ageing using hazard ratios from Cox proportional hazards models (Figure 1B). The Weibull model (i.e., power-law ageing) provided the best fit across all experimental replicates and both sexes based on Akaike Information Criterion (AIC = 111.4, R^2^ = 0.41), outperforming alternative models: Gompertz exponential ageing (ΔAIC = 7.4), logarithmic plateau ageing (ΔAIC = 11.0), logistic sigmoidal ageing (ΔAIC = 11.2), and linear ageing (ΔAIC = 17.5). The shape parameter (β = 1.07, 95% CI [0.67, 1.47]) is not distinguishable from one, so susceptibility increases approximately in proportion to age, and the data do not resolve whether the increase accelerates or decelerates over the range studied.

**Figure 1.**
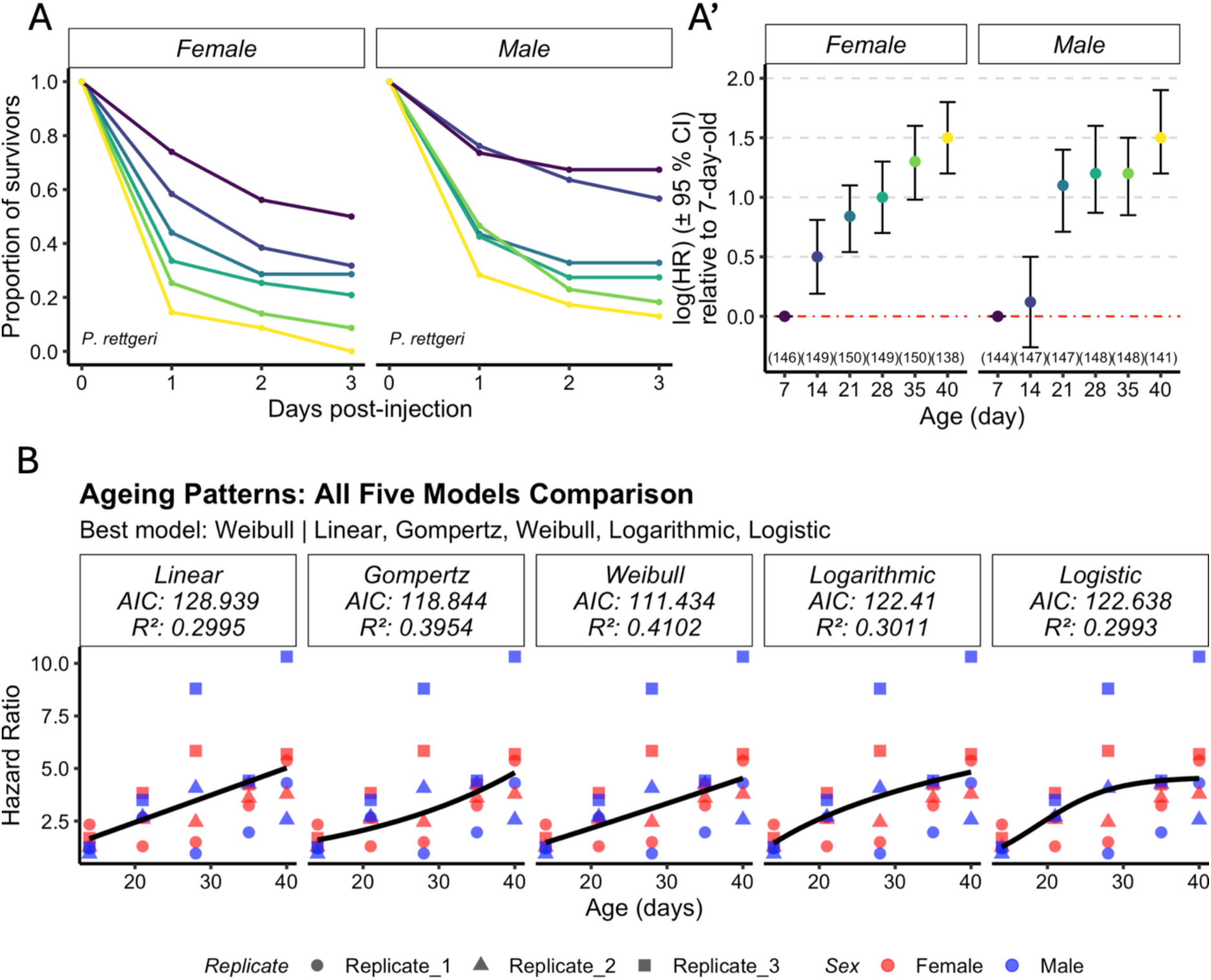
Age-related changes in susceptibility to *P. rettgeri* infection across sexes. (A–A′) Dynamics of susceptibility to systemic *P. rettgeri* infection in the RAL-821 population with age. Survival probability (A) and corresponding log hazard ratios (HR ± 95% CI) relative to baseline 7-day-old individuals (A′) were estimated for females and males infected at 7 (purple), 14 (blue), 21 (teal), 28 (dark green), 35 (light green), and 40 days of age (yellow). Colour legend of A corresponds to A’. Across all log HR plots, bars represent 95% CIs. Non-overlapping 95% CIs indicate significant differences. Sample sizes by sex and age are provided in brackets (A′). **(B) Model comparison of ageing patterns.** Using survival data from all replicates and both sexes, model comparison across five candidate ageing functions supported Weibull power-law kinetics (AIC = 111.4, β = 1.07) over Linear, Gompertz exponential, Logistic, or Logarithmic models (all ΔAIC > 7). Data from each replicate for females (red) and males (blue) are represented by different symbols.

Throughout the study, we defined 7–10-day-old individuals as “young” and 35-day-old as “old”, ensuring that despite variation between populations and infections, we can capture robust ageing phenotypes while the majority of the population remains alive.

To examine whether this increased infection susceptibility was due to inbreeding or was true on average across different genotypes, we also challenged the genetically heterogeneous host population AOx [77]. In addition to systemic infection with the Gram-negative *P. rettgeri*, we induced controlled, systemic infections with the opportunistic bacteria *Enterococcus faecalis* and *Staphylococcus aureus*. Both are Gram-positive bacteria, controlled by distinct immune mechanisms, which are in turn distinct from responses to *P. rettgeri*, which is largely controlled by a single, IMD-induced AMP, *Diptericin A* [29, 73]. Unlike in *P. rettgeri* challenge [78], the phagocytic action of hemocytes and melanisation are particularly important for surviving *S. aureus* infection [79–81], whereas the Toll/Bomanin pathway is important for surviving *E. faecalis* infection [81]. In the challenges with *E. faecalis* and *S. aureus*, we detected sexually dimorphic infection susceptibility, in which males succumbed faster than females at both young and old ages (Figure 2B’ and 2C’). Surprisingly, when infected with *P. rettgeri*, the young AOx population was not sexually dimorphic, whereas their old counterparts were (Figure 2A’). Across all challenges, infection susceptibility increased with age in both sexes (Figure 2A’’–2C’’). The overlaps between the confidence intervals suggest that the increase in susceptibility is not strongly different between sexes (Interaction in Cox model with full model; *P. rettgeri: p* = 0.025; *S. aureus*: *p* = 0.23, *E. faecalis*: p = 0.99). Finally, to test whether the age-related increase in susceptibility is also present in infections through other routes, we challenged the AOx population with the Gram-negative, entomopathogenic bacterium *Pseudomonas entomophila,* which induces a well-characterised, lethal oral infection in flies [82, 83]. Interestingly, we observed sexual dimorphism in the opposite direction to that of systemic infections: this time, males had a lower infection susceptibility compared with females [84] at both ages (Figure 2D and 2D’). Similar to what we observed in systemic infections, older individuals showed greater infection susceptibility relative to young individuals in both sexes (Figure 2D’’). These data show that with age, both sexes demonstrate increased susceptibility across a range of bacterial infections, encompassing Gram-positive and Gram-negative infections, varying levels of pathogenicity, and different routes of infection.

**Figure 2.**
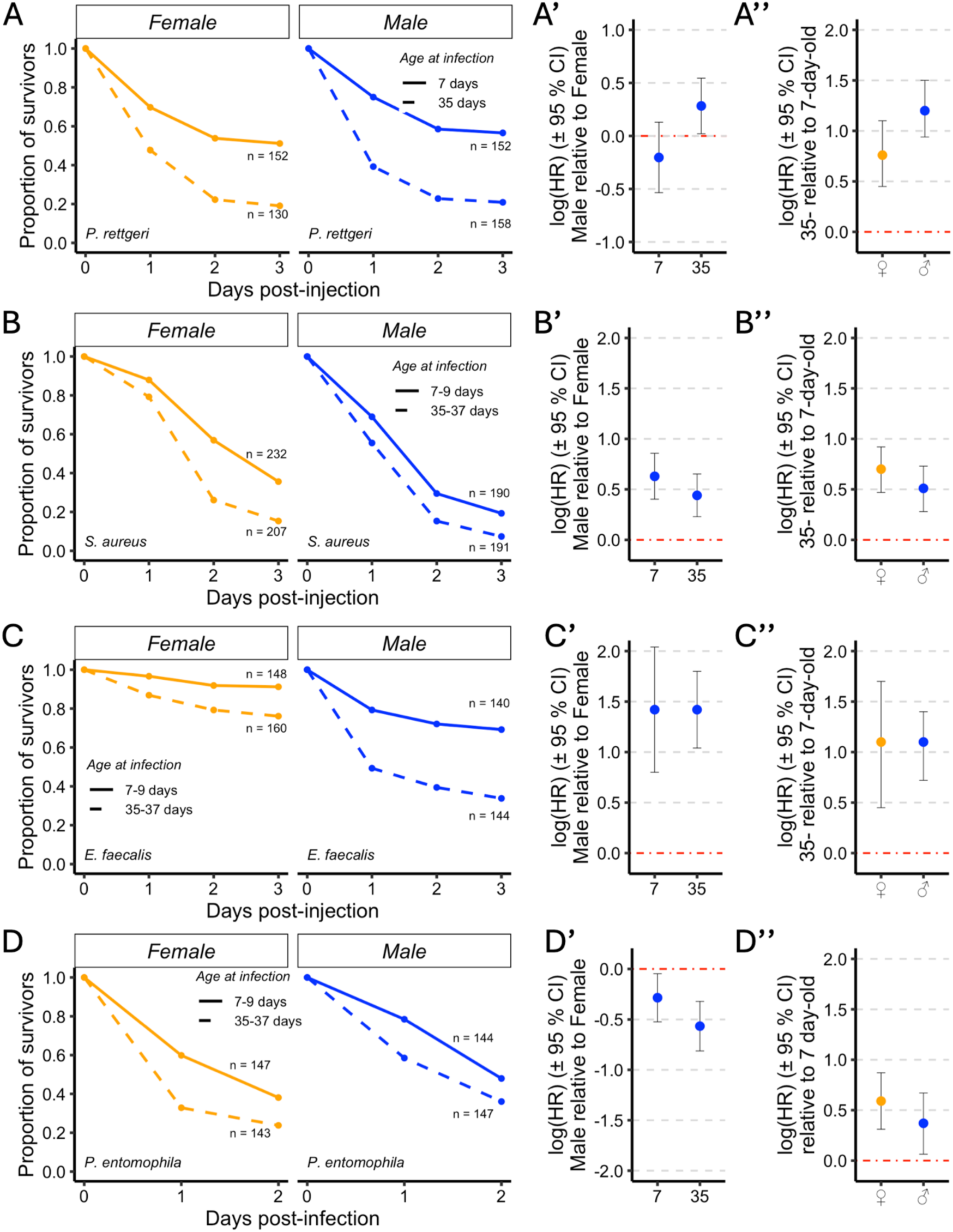
Age-related changes in susceptibility to infection across several pathogens and infection routes in both sexes. (A–D″) Changes in infection susceptibility with age are observed in both sexes across a range of pathogens and infection routes. At 7–9 (young) and 35–37 days of age (old), females and males of the AOx outbred population were exposed to four distinct bacterial challenges: systemic infections with *P. rettgeri* (A–A″), *S. aureus* (B–B″), and *E. faecalis* (C–C″), and oral infection with *P. entomophila* (D–D″). Each infection panel shows survival probability over time (A–D), log hazard ratios (± 95% CI) for infection susceptibility comparing males relative to baseline females in both age groups (A′–D′), and log hazard ratios (± 95% CI) for infection susceptibility comparing old relative to baseline young flies in both sexes (A″–D″). Across all figure panels, females and males are shown in orange and blue, respectively. Across all log HR plots, red dashed lines indicate the baseline group. Non-overlapping 95% CIs indicate significant differences. Sample sizes by sex and age are provided in the survival plots (A–D).

### Increased infection susceptibility to systemic bacterial challenge with age is widespread, but its magnitude is variable

To explore natural variation in increased infection susceptibility with age, we tested additional populations. Individuals from a panel of 24 genotypes were housed in large, mixed-sex cages, which were subsampled and subjected to *P. rettgeri* infection at 10 and 35 days old. The design of this experiment was not to characterise specific genotypes, but to generate variability and correlate infection susceptibility at different ages, lifespan, and PLUD within one large experiment. Therefore, with the exception of the block control line RAL-821, the genotypes were not replicated in different cages. With a range of approximately 5 log(HR) in both sexes (i.e., the mortality rate of the most susceptible population was about 150 times higher than the least susceptible line), we detected a high variability in infection susceptibility at young age across populations (Figure 3A). At this age, susceptibility was positively correlated between the sexes across the panel, although the ordinary least-squares estimate fell short of significance (β = 0.37, p = 0.064; Figure 3B). Because the population-level estimates differ substantially in precision, we also fitted the relationship weighting by the inverse of the estimate variance (β = 0.43, p = 0.030) and by robust MM-estimation (β = 0.42, p = 0.048), both using all populations (Supplementary Material S1, Section 3.2.2.1). The relationship was positive in all leave-one-out refits. Two populations departed sharply from this general pattern: RAL-443 and RAL-907 were among the most susceptible populations in females and among the least susceptible in males, with the sexes separated by around three log hazard units, a divergence substantially larger than in any other population (Supplementary Material S1, Section 3.2.2.2). These two populations are correspondingly the most influential on the least-squares fit, and both are retained in all analyses. Among the remaining 21 populations, in which the sexes do not diverge in direction, susceptibility is tightly correlated between males and females (β = 0.62, R²_adj_ = 0.41, r = 0.67, p = 0.001). The panel therefore comprises a majority of populations in which male and female susceptibility is closely coupled, together with a minority in which sexual dimorphism is strong enough to reverse the direction of the effect. There was also variability in infection susceptibility at old age across populations in both sexes (Figure 2 & 3C), although this was less strong – i.e., ca. 3 or 4 log(HR) than the difference in mortality rate between the extremes, which was ca. 50. However, there was no significant correlation between the susceptibility in females and in males when the individuals were old (Figure 3D).

**Figure 3.**
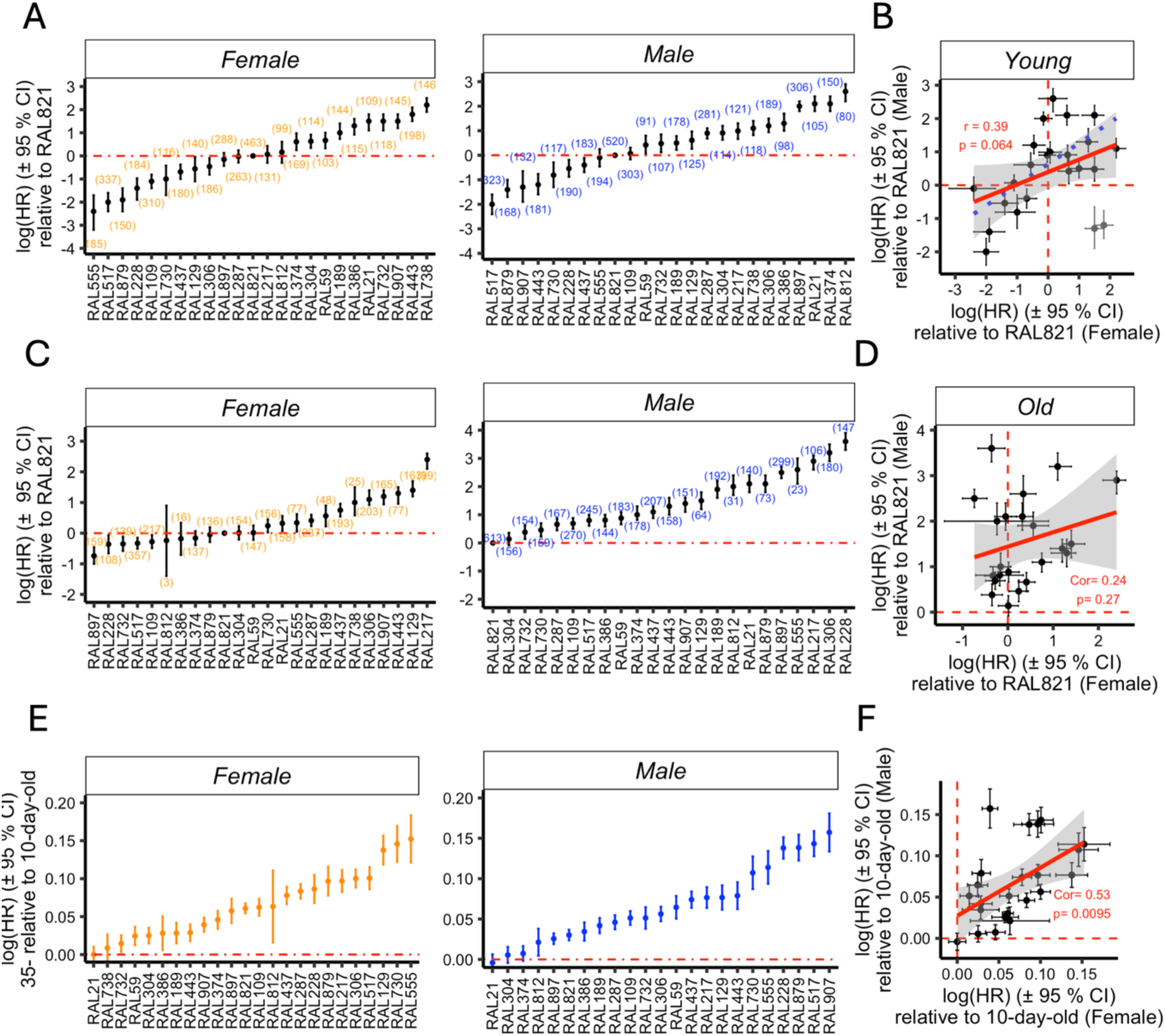
Age-related changes in susceptibility to *P. rettgeri* infection in both sexes across a panel of 24 DGRP populations. **(A)** Variation in infection susceptibility of 10-day-old (young) females and males across the panel. Log hazard ratios (± 95% CI) were estimated relative to the baseline population RAL-821. The large variation observed enables correlation of susceptibility at young age with other parameters. **(B)** Correlation between log HRs (± 95% CI) for infection susceptibility of young females and males across the panel relative to RAL-821. Line fitted by ordinary least squares to all 23 populations (β = 0.37, R²_adj_ = 0.11, p = 0.064), with the shaded band giving its 95% CI. Estimates accounting for the differing precision of the population-level values, and analyses robust to influential populations, are given in Supplementary Material S1, Section 3.2.2.1. Grey points mark RAL-443 and RAL-907, the two most strongly sexually dimorphic populations in the panel; both are included in the fit. The blue dotted line shows the fit to the remaining 21 populations (β = 0.62, R²_adj_ = 0.41, p = 0.001), in which the sexes do not diverge in direction. **(C)** Variation in infection susceptibility of 35-day-old (old) females and males across the panel. Log HRs (± 95% CI) were estimated relative to the baseline population RAL-821. The large variation observed enables correlation of susceptibility at old age with other parameters. **(D)** Correlation between log HRs (± 95% CI) for infection susceptibility of old females and males across the panel relative to RAL-821. No significant correlation was found (β = 0.32, R^2^_adj_ = 0.01, r = 0.24, p = 0.27). **(E)** Age-related differences in infection susceptibility of females and males across the panel. Log HRs (± 95% CI) were estimated for old females and males within each population relative to their respective young baselines. Old individuals were predominantly more susceptible than young ones, indicating that age-related increases in susceptibility are widespread. A small number of populations showed no detectable increase; these were among the most susceptible at 10 days of age, but were not at any upper limit of measurable susceptibility (see Results and Supplementary Material S1, Section 3.2.5). **(F)** Age-related changes in infection susceptibility are positively correlated between females and males. Correlation between log HRs (± 95% CI) for infection susceptibility of old females and males within each population relative to their respective young baselines (β = 0.58, R^2^_adj_ = 0.26, r = 0.53, p = 0.01). Across all log HR plots (A, C, E), females and males are shown in orange and blue, respectively. Across all log HR correlation plots (B, D, F), dots represent estimates from the models, and solid red lines show regression fits with grey shading representing 95% CIs. Red dashed lines indicate baseline group estimates, and bars denote 95% CIs. Non-overlapping 95% CIs indicate significant differences. Sample sizes by sex and population are shown in brackets (A, C, E). Correlation coefficients and p-values are provided in each plot (B, D, F). RAL-821 was excluded from the correlation calculations in (B) and (D), where all other populations are estimated relative to it and its inclusion would violate the assumption of independence. In (F), where each population is compared with its own young baseline, RAL-821 has an estimated value like every other population and is included.

We then compared infection susceptibility between flies of each sex sampled from the same cage at young and old ages. This demonstrated that the increased susceptibility in aged individuals across the panel was widespread (Figure 3E). Both sexes of RAL-21 and males of RAL-304 and RAL-374 were an exception, and did not show a detectable increase in susceptibility over age. These populations were among the most susceptible at 10 days of age (Supplementary Material S1, Section 3.1), but this does not reflect an upper limit on measurable susceptibility: at 35 days their log hazard ratios ranged from 0.14 to 2.1, well below the maximum of 3.6 observed across the panel. Rather, they lie at one end of a continuum. Susceptibility at 35 days was largely unrelated to susceptibility at 10 days, with slopes of old on young of 0.11 in both sexes, far below the slope of one expected if the age-related increase were independent of baseline susceptibility (p ≤ 3 × 10⁻⁴; Supplementary Material S1, Section 3.2.5). Populations that were already susceptible when young therefore gained least with age, and the ordering established early in life did not persist. Simulations confirmed that this convergence cannot be produced by measurement error or regression to the mean alone. The clearest illustration is given by RAL-443 and RAL-907: the sexes are separated by 3.0 and 2.8 log hazard units at 10 days, but by 0.0 and 0.2 at 35 days, so the strong dimorphism present early in life is absent at old age. Despite the lack of correlation between sexes for infection susceptibility at old age, there was a positive correlation in the increase in susceptibility over ageing (Figure 3F).

### Increase in infection susceptibility occurs through both loss of resistance and loss of tolerance and varies by population

We sought to gain insight into the strategies associated with age-related changes to infection susceptibility. We utilised an approach based on our recent combined theoretical modelling and empirical testing in *D. melanogaster* [42]. The premise of this approach is that <u>P</u>athogen <u>L</u>oad <u>U</u>pon <u>D</u>eath (PLUD) – a tolerance metric [45] – and mortality measures like hazard ratios (HR), when estimated across multiple populations, can be used to predict a relationship between PLUD and infection susceptibility. A negative relationship between PLUD and HR implicates defects to tolerance in driving increased susceptibility, while a positive relationship implicates decreased resistance [42].

Survival data from *P. rettgeri* infections of young and aged individuals across a panel of 24 populations were used to obtain estimates for age-related changes in infection susceptibility (Figure 3). To quantify PLUD across the panel, individuals were sampled within 30 minutes of their death between ∼16–24 hours post-infection. PLUD estimates were variable across lines and sexes (Figure 4; Supplementary Material S1, Section 4.1). Age was found to increase or reduce the PLUD in a population- and sex-specific manner (Figure 4A). We then combined the estimates for PLUD and age-related changes in infection susceptibility from RAL populations to determine the main reason underpinning the latter. We had PLUD estimates for 19 populations for males and 23 for females. We found that, across the panel, age-related susceptibility was associated with reductions in both resistance (e.g., RAL-821 and RAL-907) and tolerance (e.g., RAL-228 and RAL-555) (Figure 4A).

**Figure 4.**
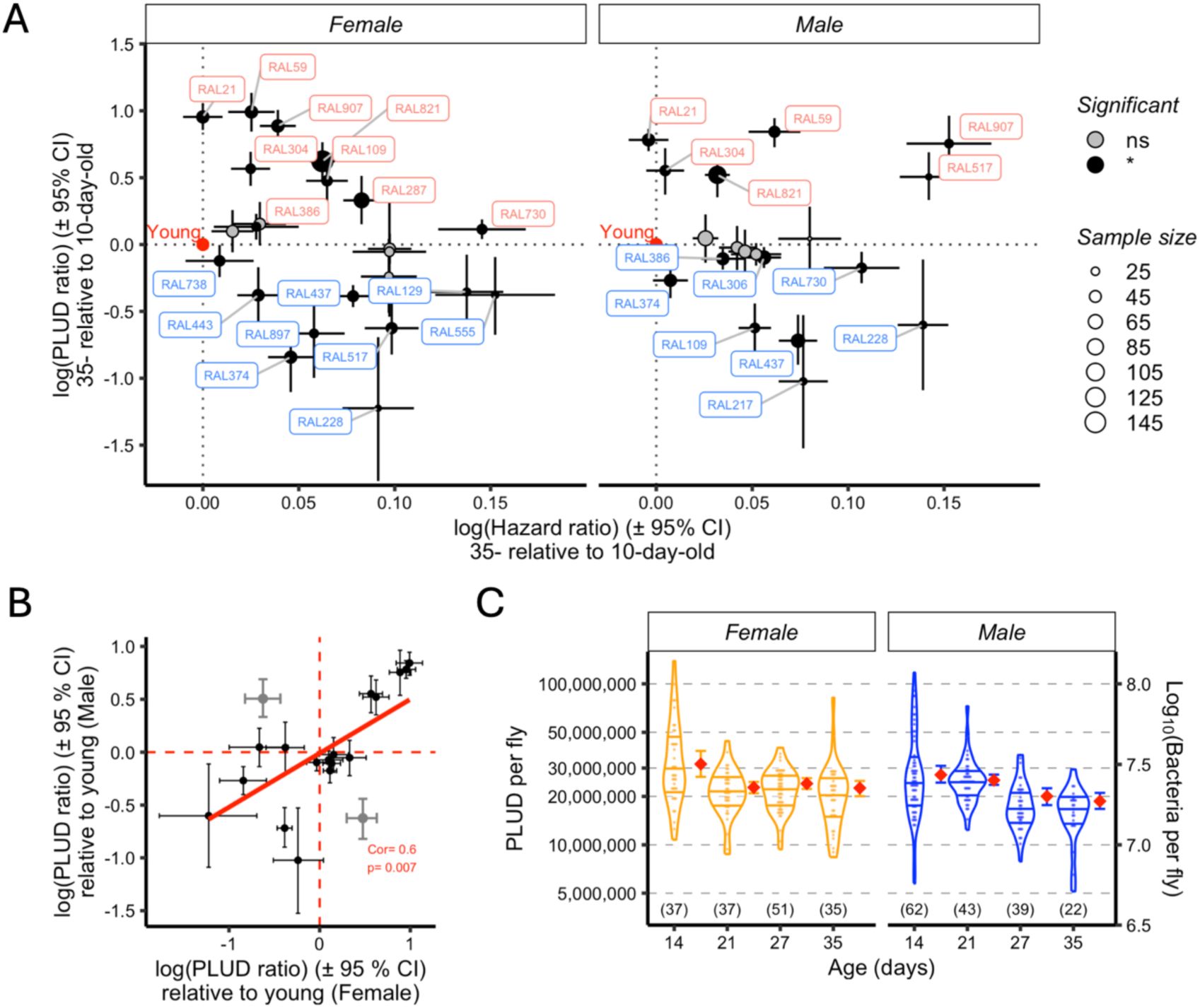
Harnessing Pathogen Load Upon Death (PLUD) to investigate age-related changes in infection susceptibility. **(A)** Relative contribution of resistance and disease tolerance to age-related increases in infection susceptibility. PLUD and susceptibility (HR) of old relative to their young counterparts across 23 female and 19 male DGRP populations. The log PLUD ratio (± 95% CI) and log (HR) (± 95% CI) together indicate the relative contribution of resistance and tolerance to infection susceptibility (Duneau *et al.*, 2025). Both loss of tolerance and loss of resistance can explain increased infection susceptibility with age. Populations that become primarily tolerance-deficient with age show low PLUD ratios (blue labels), while those primarily becoming resistance-deficient populations show high PLUD ratios (salmon labels). Filled circles indicate significant age effects (95% CI excludes zero). Point size reflects sample size. **(B)** Age-related changes in PLUD are correlated between the sexes. Correlation between female and male log PLUD ratios (± 95% CI) across 18 DGRP populations relative to the baseline population RAL-821, suggesting that in most populations both sexes die for the same underlying reason (β = 0.67, R²_adj_ = 0.39). All 18 populations shown are included in the correlation and in the fitted regression. RAL-517 and RAL-109 are highlighted in grey because the sexes change in opposite directions in these two populations, they are not excluded from any analysis. RAL-129 and RAL-879 are absent because male sample sizes were insufficient to estimate PLUD. RAL-821 is excluded from the correlation, as all other populations are estimated relative to this reference and its inclusion would violate the assumption of independence. **(C)** Age-related changes in PLUD in the population RAL-228 where increase in susceptibility is mainly driven by a decrease in tolerance. With age, the PLUD changes in the same direction, indicating that populations dying mainly from a decrease in tolerance when old were not dying from deficient resistance when younger. PLUD is displayed on a log_10_ scale (right y-axis) with corresponding natural scale values (left y-axis) to better convey the absolute magnitude of bacterial load differences, which span millions of bacteria. Violin plots represent PLUD values for each individual and the distribution of the samples, while red diamonds represent the model-estimated PLUD (see Methods), which is weighted by the precision of each individual estimate and is therefore not the arithmetic mean of the plotted values. Females and males are shown in orange and blue, respectively, and sample sizes are provided in brackets (C). Across all panels, bars represent 95% CIs. Non-overlapping 95% CIs indicate significant differences.

There was a strong correlation between sexes (Figure 4B), although we detected scenarios where, within a population, males showed reduced resistance over ageing while females showed reduced tolerance (e.g., RAL-517); the opposite was also true (e.g., RAL-109). These findings suggest that age-related infection susceptibility is often associated with the decline of non-immune functions, and that the underlying cause is usually the same for both sexes.

### Dynamics of PLUD over ageing

To characterise the dynamics of the reduced tolerance over age, we measured PLUD at weekly intervals in the tolerance-deficient genotype RAL-228 (Figure 4C). We hypothesised that a reduction in resistance could first impact infection susceptibility, followed by an accumulation of declines in non-immune functions that would have an impact on tolerance at a later age, but this was not what we observed. The PLUD demonstrated a step change in both sexes, dropping and remaining low up to 35 days of age. These data suggest that once ageing affects mechanisms that support disease tolerance in early middle age, the PLUD decreases and remains low.

### Increase in infection susceptibility is an early-arising ageing phenotype

Since the magnitude of the age-related increase in susceptibility varied significantly, we were able to correlate it with other parameters to better understand it. We sought to directly measure the relationship between both early- and late-life infection susceptibility and lifespan of unchallenged individuals. Lifespan and *P. rettgeri* susceptibility data were collected in parallel across a panel of 12 DGRP lines, with 4 lines sampled in each experimental block (except Block 2, which included 3 lines) (Figures 3 and 5, respectively, and Supplementary Material S1, Section 5.1 for lifespan curves). Lifespan was highly repeatable for RAL-821 across experiments in both sexes, and although Block 1 lived less long on average, this confirms that the variability across populations is not only due to experimental or environmental variation (Supplementary Material S1, Section 5.2). As expected, there was extensive phenotypic variation in lifespan across populations (Figure 5A), which was strongly correlated between sexes (Figure 5B, linear model with Block covariate: β = 1.09, p = 0.03; raw correlation r = 0.90, p < 0.001). To test whether this correlation could have been driven by Block 1 – despite using Block as a covariate – we repeated the analysis excluding Block 1: the correlation remains substantial (r = 0.6, p = 0.1, Supplementary Material S1, Section 5.4.2), though statistical power is reduced with fewer populations (n = 8 vs. n = 11). The effect size of r = 0.60 represents a strong correlation, supporting that this reflects a genuine biological pattern.

**Figure 5.**
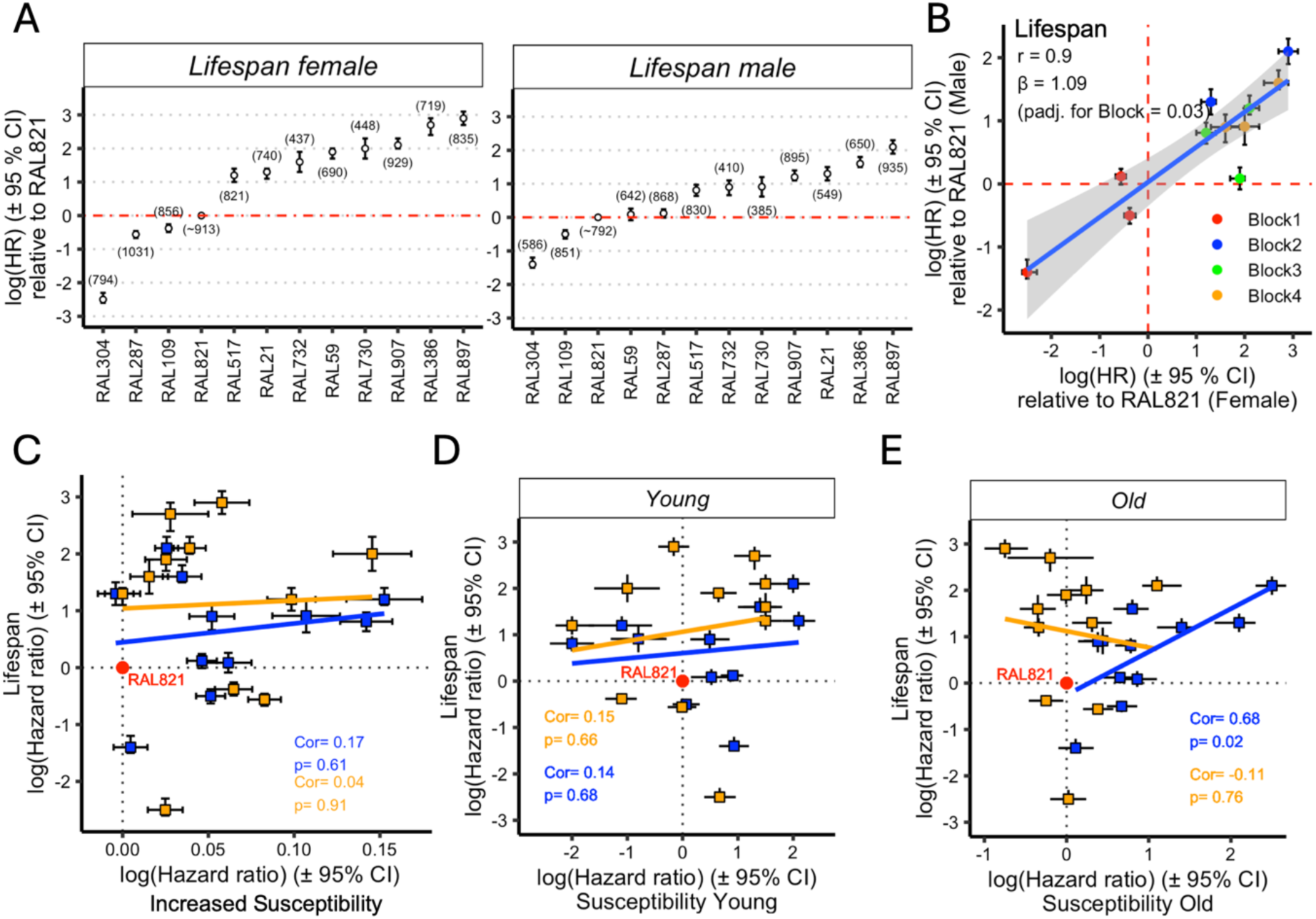
Correlation between infection susceptibility and lifespan in controlled conditions. **(A)** Variation in female and male lifespan across a panel of 11 DGRP populations relative to the baseline population RAL-821. Log hazard ratios (HR ± 95% CI) for lifespan of females and males across the panel relative to the baseline population RAL-821. **(B)** Female and male lifespan are strongly correlated (linear model with Block covariate: β = 1.09, p = 0.03; raw correlation r = 0.9, p < 0.001). Each point represents a population, with log (HR) (± 95% CI) calculated relative to RAL-821 within each block. **(C)** Infection susceptibility at young age does not predict lifespan. **(D)** Increased susceptibility at old age is correlated with shorter lifespan in males (β = 0.5, R^2^_adj_ = 0.41) but not in females (β = –0.3, R^2^_adj_ = –0.1). **(E)** The age-related increase in infection susceptibility does not predict lifespan. Across all figure panels, within each sex and experimental block, estimates are relative to the baseline population RAL-821. Baseline estimates are indicated by red dashed lines (A–B) or black dotted lines (C–E). Female and male estimates are shown in orange and blue, respectively (C–E), and each experimental block is represented by a distinct colour (B). Sample sizes by sex and age are provided in brackets (A). Non-overlapping 95% CIs indicate significant differences. Correlation coefficients and p-values are shown in each plot (B–E). Solid lines represent regression fits (B–E), with grey shading indicating their 95% CIs (B). Survival curves for all populations are shown in Supplementary Material S1, Section 5.1. RAL-821 was excluded from correlation calculations as all other lines are estimated relative to this reference population, which would violate the assumption of independence.

Notably, virtually all lines show a strong increase in infection susceptibility by 35 days, which is before the median lifespan for most populations (Supplementary Material S1, Section 3.1). We have granular data for the genotype RAL-821: in addition to weekly *P. rettgeri* infection data (Figure 1), we have produced high-powered lifespan data, as it was used as a block control, giving four independent replicates with over 700 individuals per sex in each. In this genotype, we can detect a strong increase in infection susceptibility in both sexes by 21 days, an age where the probability to die naturally is close to zero and approximately one-third of the average median lifespan for the line (i.e., ∼60 days). These data suggest that increased susceptibility to infection can be an early-arising ageing phenotype, with potential to drive systemic ageing [67, 85].

### Relationship between early and late-life infection susceptibility and lifespan

To relate infection susceptibility in early- and late-life to lifespan, we fitted a linear model. This used sex- and population-specific log hazard ratio (HR) estimates. These HR values quantify relative differences in both lifespan and infection susceptibility across populations. The RAL-821 population was not included in those analyses as it was used to rank the other populations, and therefore is not an independent observation. This restricted our analyses to 11 populations. The age-related increase in infection susceptibility – measured as the relative difference between old and young individuals – did not correlate with lifespan in either sex (Figure 5C). Early-life infection susceptibility and lifespan were not significantly associated in either sex (Figure 5D). However, there was a strong positive relationship between late-life susceptibility and lifespan, but only in males (Figure 5E).

## Discussion

Infections challenge hosts in a variety of ways, engaging different features of the host response, and requiring costly resources to mount defence and repair mechanisms. Both the ability to mount an efficient immune response (i.e., resistance) and to tolerate the consequences of the infection (i.e., tolerance) are assumed to decrease with age, but the relative impact of their decline on the susceptibility of older individuals can be difficult to untangle. Variation within populations in the ability to maintain immune competence over age, well-documented in humans [5, 6], suggests that individuals’ immune systems may fail in different ways and at different times. It is particularly challenging to know when the decline in non-immune mechanisms have an impact on host susceptibility to infection. Furthermore, the chronology of physiological declines and their implications could vary depending on genotype, including sex, on environmental factors, or on their interaction. Analysing pathogen load – i.e. asking the pathogen to report on host responses – offers a way to uncover chronology of increasing susceptibility and the relative contribution of declining resistance and tolerance [42]. Using *D. melanogaster*, we have shown that the increase in infection susceptibility over ageing can start early in life and is widespread across a range of bacterial pathogens and routes of infection, and occurs in both sexes. Leveraging the variation generated by infecting both sexes of multiple host genotypes with a single bacterial pathogen, we showed that susceptibility when young and its decline correlate between sexes while susceptibility when old does not. We also showed that the increased susceptibility can be attributed to a decline in resistance, but most frequently to a decline in tolerance. By correlating infection susceptibility at different ages with lifespan, we demonstrated that immune competence at old age predicts longevity in males only, regardless of the existence or direction of sex bias in lifespan.

### Increase in infection susceptibility as an early arising ageing phenotype

A standing question in the field of immune ageing is whether increased infection susceptibility arises only in late life, or begins to manifest earlier. Increase in susceptibility to infections and ageing appear to be linked, with cause and consequence difficult to tease apart [86]. Observational studies in humans suggest that immune function may begin to decline surprisingly early, with measurable changes from adolescence [87–89] and distinct physiological transitions identified by the third decade of life [90, 91]. However, direct experimental evidence that increased susceptibility precedes natural death has been inconclusive [92]. Although age-associated infection susceptibility has been described in *D. melanogaster* [66, 67], studies with sufficient longitudinal resolution to detect the early onset of immune decline are rare [93].

Here, we provide such longitudinal resolution and empirical evidence that increased infection susceptibility is an early-arising ageing phenotype in *D. melanogaster*. We show that susceptibility to *P. rettgeri* increases significantly between 14 and 21 days of age, about one-third of the median lifespan (i.e., ∼60 days) in the focal genotype RAL-821. Notably, this decline in immune competence occurs while flies are still within their peak reproductive period and actively engaging in mating [94, 95]. This indicates that age-related increase in susceptibility can emerge well within the physiological “prime” of life, consistent with evolutionary theories proposing that ageing evolves in response to patterns of external mortality and reproductive investment [96–98]. In the wild, where *D. melanogaster* face high extrinsic mortality from environmental hazards such as predation [99–101], selection for maintaining immune function in later life is likely weak. Consequently, resources may be preferentially allocated to early reproduction over long-term somatic maintenance, leading to immune decline even while individuals remain reproductively active [102, 103]. Our findings highlight the importance of considering early-life immune decline in ageing models, not just late-life immune failure.

### Populations converge in susceptibility with age

Understanding the rate and pattern of ageing of biological processes is a fundamental question with important implications for health and longevity. We detected a correlation between male and female infection susceptibility when young, suggesting that the sexes shared similar baseline susceptibility, strongly influenced by development and genotype. Furthermore, the magnitude of age-related increase in susceptibility was also correlated between sexes. However, somewhat counter-intuitively, the correlation in susceptibility broke down at old age. This breakdown could be attributed to environmental factors that accumulate differentially during ageing, such as lifestyle differences or varying environmental stresses. However, our study was performed under standardised and constant laboratory conditions where sexes were mixed, suggesting that environmental heterogeneity cannot fully explain this pattern, unless considering an interaction between genotype – including sex – and environment. Instead, our findings could point to a fundamental characteristic of biological ageing processes.

A longstanding question in ageing research is how quickly biological processes change with age. Since Gompertz described mortality as increasing exponentially with age [104], it has become clear that each unit of time adds a constant to log-mortality rate, resulting in multiplicative increases in the chances to die. However, not all processes necessarily age with the same dynamic, so we tested whether infection susceptibility follows different ageing patterns: Linear (i.e., constant ageing), Gompertz (i.e., exponential ageing), Weibull (i.e., power-law ageing), Logarithmic (i.e., ageing with asymptotic plateau), or Logistic (i.e., ageing with threshold). Our data support a Weibull framework with a shape parameter that is not distinguishable from one (β = 1.07, 95% CI [0.67, 1.47]), operating in log-space as log-susceptibility = baseline + rate × log(age). While the fitted curves appeared visually similar to the linear model in linear-space (Figure 1B), the substantial AIC difference (ΔAIC = 17.5) reflects fundamentally different underlying assumptions: the Weibull model assumes multiplicative ageing effects (i.e., proportional increases), whereas the linear model assumes additive effects (i.e., constant increases). The confidence interval on the shape parameter is wide, and we therefore make no claim that susceptibility either accelerates or decelerates with age. What the comparison establishes is that the increase is better described as a power function of age than as the exponential function that describes overall mortality risk, so susceptibility does not accelerate into late life in the way a Gompertz process would predict.

Our analysis combined both sexes to maintain sufficient statistical power for comparing complex non-linear ageing models. While we assumed a universal ageing pattern across sexes, future work with larger sample sizes could explicitly test whether males and females follow different ageing trajectories. However, the strong correlations we observe in baseline susceptibility and age-related changes suggest that any sex-specific differences in ageing dynamics are likely to be subtle. Males and females show correlated susceptibility early in life and in age-related changes (Figure 3B and 3F), while the correlation is weaker at old age (Figure 3D). This is accompanied by a marked convergence between populations: susceptibility at 35 days is almost independent of susceptibility at 10 days, so the ordering established early in life does not persist and correspondingly less shared variation remains for the sexes to correlate on. What determines susceptibility at old age is therefore largely not a continuation of what determines it when young.

These findings indicate that infection susceptibility over ageing follows Weibull power-law dynamics commonly observed for individual biological traits, distinct from the Gompertz exponential pattern typically seen in overall mortality risk. This confirms that trait-specific ageing may follow different mathematical patterns than organismal-level mortality [105], with biological damage in specific systems following power-law dynamics that operate somewhat independently of initial population differences.

### Sexual dimorphism in ageing processes

We detected a correlation between the sexes in the magnitude of the age-related increase in susceptibility (Figure 3F), suggesting that both sexes suffer a decline in immune function. Strikingly, when we examined the covariance of susceptibility at old age with lifespan across populations, the risk to die from infection predicted lifespan, but only in males, regardless of the existence or direction of sex bias in longevity. In other words, this association is not a consequence of males being the shorter-lived sex. Populations in our panel differ in which sex lives longer, and in some the bias is reversed (Figure 5A), yet the relationship between late-life susceptibility and lifespan is confined to males throughout. Had the association simply tracked whichever sex was more frail, we would expect it to follow the direction of the sex bias rather than the sex itself. This suggests that increased susceptibility to infection is a driving risk factor for mortality in males, whilst other features of ageing, such as the well-documented decline in intestinal integrity [63, 106–109], are more important to determine length of life in females [65, 110]. Our data add to burgeoning evidence across taxa that males suffer more from reductions in immune function over age [111–115], which is predicted by evolutionary theories of ageing that posit females will derive a greater increase in fitness by investing for longer in immune competence, compared to males who primarily direct investment into competitive features of reproduction [116–119].

### Increase in infection susceptibility can frequently be due to a decline in non-immune functions

We found that increased infection susceptibility resulted from declines in both direct immune resistance mechanisms and functions that support the ability to tolerate the infection. It has been shown that the comparison of pathogen load upon death (PLUD) between groups with different susceptibilities can tell whether the difference is predominantly due to defects in resistance or tolerance [42]. We used this method to compare the PLUD between young and old individuals across 23 populations. Our data identify several populations where increased mortality was linked with a decline in resistance, characterised by higher infection susceptibility and a higher PLUD. Examples of functional declines in resistance mechanisms over ageing have not been well-described, with a few exceptions, such as the reduction in actively phagocytosing hemocytes in circulation [61], together with deficits in the processing capacity within phagocytic vesicles [62], and changes to AMPs expressed upon infection [60]. Yet, how these changes directly – and collectively – contribute to age-associated susceptibility remain to be determined.

We showed that a slim majority of populations had a lower PLUD, and therefore were more susceptible at old age mainly due to a decrease in their ability to sustain the infection. Some aspects or physiologies of tolerance overlap with that of the molecular biology of ageing. Tolerance is linked to intracellular functions such as endoplasmic reticulum maintenance and the protein-folding response [29, 41], and systemic functions such as metabolic flexibility, and the dysregulation of these processes are hallmarks of ageing [120]. Additionally, tolerance is mediated through regulation of immune signalling [121–123], known to be dysregulated with age [55, 56, 88, 124]. As such, we might expect age to negatively affect tolerance. However, only a handful of research mechanistically assesses age-related immunity in *D. melanogaster* and elsewhere, especially in terms of tolerance [54, 125]. Increased susceptibility to *E. coli* with age was found to be unrelated to changes in resistance, and suggests a reduction in tolerance with age [126]. Age-related susceptibility to Flock House virus was shown to be tolerance-dependent [54], and dysregulation of AMPs upon bacterial infection can damage tissue, impacting function [60]. Our results support the need to pursue further research into tolerance-dependent, increased susceptibility to infections over age.

There are concerted efforts to maintain immune responses to acute infections in elderly humans [127, 128], and therapeutic targeting of the immune system has been posed as a promising anti-ageing intervention [129–131]. Our data strengthens the argument that individuals do not age uniformly, that population variation –including sex – can determine routes to age-related increase in infection susceptibility, and that susceptibility can increase due to causes which are not directly linked to immune decline *per se*. This will be an important consideration for developing strategies to limit susceptibility in older individuals. We have shown that, by interrogating pathogen load alongside infection susceptibility, the combined effects of declining systems can be measured and attributed to failures in resistance or tolerance. Our findings establish that further work across systems should not only focus on *bona fide* immune dysregulation of resistance mechanisms within immune tissues, but should also concentrate on the dysregulation of non-immune functions, and on how they impact the ability to control infections. As such, studies of age-related infection susceptibility should not use the term “immunosenescence” without demonstrating that increased susceptibility is directly linked to a decline in *bona fide* immune functions.

## Conclusions

This study has shown that increased infection susceptibility is an early-arising ageing phenotype in *Drosophila.* It occurs in both sexes across a range of pathogens and routes of infection, yet drives mortality in males only, regardless of the existence or direction of sex bias in longevity. By leveraging natural genetic variation across a panel of isogenic lines, we show that increased infection susceptibility most frequently reflects a failure of disease tolerance rather than a failure of canonical immune resistance mechanisms, although both can occur. We therefore argue that the term “immunosenescence” should be limited to cases where increased susceptibility is demonstrably linked to dysregulation of *bona fide* immune resistance mechanisms. Fully understanding age-related loss of pathogen control will require explicit consideration of population variation, host sex, and the broader ageing of non-immune functions that shape the host’s ability to tolerate infection.

## Methods

### D. melanogaster stocks and husbandry

Flies were maintained at 25 °C in 60–70% humidity on a 12:12 light:dark cycle. Diverse infections were tested in a genetically heterogenous line, the Ashworth Outcrossed (AOx) population – a large, lab-adapted outbred *D. melanogaster* population derived from the DGRP panel [69]. The AOx was originally established by setting up 100 crosses of 113 DGRP lines [77] and has since been maintained as an outbred population on a 14-day generation cycle, with controlled larval density, and a census size of between 3,000 and 4,000 adults in each generation [132]. To generate variability and correlate infection susceptibility at different ages, lifespan, and PLUD within one large experiment, 24 single isogenic lines from the DGRP [69] were obtained from the Bloomington Drosophila Stock Center (NIH P40OD018537). Associations between lifespan and susceptibility were assessed in 12 DGRP lines: RAL-21, RAL-59, RAL-109, RAL-287, RAL-304, RAL-386, RAL-517, RAL-730, RAL-732, RAL-821, RAL-897, RAL-907. These lines were also used to assess age-related changes to infection susceptibility and PLUD, in addition to 12 other genotypes: RAL-129, RAL-189, RAL-217, RAL-228, RAL-306, RAL-374, RAL-437, RAL-443, RAL-555, RAL-738, RAL-812, and RAL-879. All lines were expanded in a standardised, density-controlled manner in which an average of 400 eggs were dosed into bottles containing standard Lewis food [133], and this process was repeated for a minimum of four generations to control for transgenerational effects. To replicate natural conditions and minimise the impact of social isolation on immunity and lifespan [134, 135], all experimental flies were maintained in mixed-sex, plastic cages. For infections presented in Figures 1 and 2, these measured 15 cm × 8 cm × 15.5 cm and were fitted with two standard Lewis food vials, which were replaced every other day. Cages were set up when flies were 0–2 days old, with an average density of approximately 150 flies per cage, and 100 flies per cage after infection. All experiments were conducted across at least three replicates.

### Survival to diverse bacterial infections

To assess age-related changes in infection susceptibility, we parameterised our infections using a single isoline (RAL-821) infected with the Gram-negative bacteria *Providencia rettgeri* [74]. We infected individuals at 7, 14, 21, 28, 35 and 40 days of age to determine the change in susceptibility over time. Based on this *P. rettgeri* infection time course, we termed 7–10 days old as “young”, and 35 days old as “old”, with the latter term used in relative contrast to young. We selected this aged timepoint as it allows for meaningful comparisons across populations: flies exhibit a clear increase in infection susceptibility, yet the majority of the population remains alive, therefore minimising selection bias (as confirmed by our lifespan data, with >90% of flies alive at 35 days of age).

#### Systemic infections

Between 3–4 days prior to infection, LB agar plates were streaked with either *P. rettgeri* (Dmel strain)*, S. aureus* (PIG1 strain) *or E. faecalis* (OG1RF strain) from frozen 25% glycerol stocks and incubated overnight at 37 °C. A single colony was suspended in LB broth and incubated overnight at 37 °C (24 hours for *E. faecalis*). The resulting culture was pelleted and resuspended in PBS to an optical density at 600 nm (OD_600_) of 0.1. Flies were anaesthetised with CO_2_ and injected in the abdomen with 23 nL of bacterial suspension using a Nanoject III microinjector (Drummond) [136]. Flies were transferred to mixed-sex cages, and mortality was recorded twice daily for three days. This period captures the acute infection phase characteristic of the bacterial species used, during which mortality differences are most pronounced and directly reflect the host’s ability to control bacterial proliferation [45]. This timeframe also avoids introducing additional sources of mortality unrelated to infection susceptibility (e.g., age-related mortality, especially in older flies), which would confound quantification of infection-induced mortality.

#### Oral infection with P. entomophila

1% skim milk LB agar plates were streaked from frozen 25% glycerol stocks and incubated at 30 °C until single colonies were visible. A single protease-positive colony – identified by the presence of a surrounding proteolytic ring [137] – was suspended in 10 mL of LB broth. After 5 hours, 400 µL of the resulting culture was divided across two large flasks of 200 mL LB broth, and expanded over a further 24-hour period. Bacterial suspensions were pelleted by centrifugation at 4 °C and 2,500 × g for 15 minutes. The pellet was resuspended in 5% sucrose water to an OD_600_ of 100 [137]. 150 µL of either 5% sucrose water or bacterial sucrose suspension was applied to a Whatman filter paper, placed on top of Lewis food in a vial, and left to dry for approximately 30 minutes prior to infection. Flies were starved in empty vials for 3 hours at 25 °C before being transferred to infection vials, where they were allowed to feed overnight for 16 hours. The following day, they were moved to fresh food vials, and mortality was recorded daily for two days, a period chosen to capture the acute phase of this infection, following the same rationale described above.

### DGRP panel infection susceptibility and lifespan assay

For experiments in Figures 3, 4 and 5, flies of each line were housed in individual 15.5 × 39.5 × 25.5 cm plastic cages containing three 90 mL Petri dishes with Lewis food, which were replaced every other day. The cages were set up when flies were between 0–2 days old, and flies were sorted under CO_2_-induced anaesthesia to allow for the initial density of males and females to be estimated (i.e., on average 718 and 796 per cage, respectively). The experiment consisted of 4 blocks. Each experimental block consisted of 7 populations, including a block control, the line RAL-821. Infection susceptibility, bacterial load and lifespan (for half of the lines) were measured for each population in a single experiment. In each block, lifespan was measured in 4 out of the 7 populations (3 in Block 2). Because each population was housed in a single cage, population identity and cage are confounded in this design, and the differences we report between populations reflect the combined contribution of genotype and cage environment. RAL-821 was included in every block to quantify and adjust for variation between experiments. All traits for a given population (lifespan, early- and late-life susceptibility, and PLUD) were measured on individuals sampled from the same cage, so the correlations we report among these traits are unaffected by this confound. The differences between populations therefore reflect the combined contribution of genotype and cage environment, and we do not attribute them to genotype alone.

#### Lifespan analysis

Deaths in each cage were scored every other day through the removal of corpses from the cages, and their subsequent sexing. Flies used for infections were removed from cages and censored. Lifespan was recorded until all flies had died.

#### Infection susceptibility analysis

The evening preceding the infection, flies were collected and housed in bottles with a balanced sex ratio. *P. rettgeri* was prepared as described above. 23 nL of bacterial suspension was injected into the fly abdomen, corresponding to a dose of approximately 3,500 viable bacteria per fly. Flies were sedated with CO_2_ during the injection procedure, and then added to individual 15 × 15 × 16 cm plastic cages with 90 mL Lewis food. Deaths were scored roughly every 30 minutes from approximately 12–24 hours post-infection, and 2–3 times from 36–48 hours post-infection.

#### Measuring Pathogen Load Upon Death (PLUD)

To assess the strategies associated with age-related changes in infection susceptibility, PLUD was measured for each sex of each line. This was done by monitoring and collecting flies within 30 minutes of death between 14–24 hours post-infection, with death characterised by a lack of movement and responsiveness. The corresponding infection susceptibility assays (i.e., survival post-infection) were monitored in parallel until the end of the 48-hour period. PLUD represents the bacterial load threshold at which hosts succumb to infection. This metric has been shown to be remarkably stable over time post-infection and largely independent of initial inoculum dose, making it a robust measure that reflects the integrated contributions of both resistance (i.e., the ability to control bacterial proliferation) and disease tolerance (i.e., the ability to sustain bacterial burden) to infection susceptibility [42, 45]. Flies were placed individually into 1.5 mL microcentrifuge tubes containing 250 µL of PBS and two 3 mm glass beads before homogenisation in a TissueLyser II (Qiagen) at 25 Hz for 30 seconds. Tubes were kept at 4 °C for plating the following day. Duplicates of samples were serially diluted 1:10 in PBS thrice, and the first two of these dilutions were discarded. The suspensions were subsequently serially diluted 1:3 in PBS with the final dilution being 1:2,187,000. 5 µL of each suspension was plated onto selective LB agar plates supplemented with erythromycin at a concentration of 5 µg/mL, to which *P. rettgeri* is naturally resistant [138]. Plates were left overnight at room temperature and then placed in an incubator at 37 °C and monitored frequently over several hours. This procedure allows for counting small and non-overlapping colonies.

### Statistical analysis

All analyses were conducted in R using RStudio.

#### Computation of estimated Hazard Ratio (HR) and 95 % confidence intervals

Survival analyses were conducted using the function *coxph* from the package *survival.* The confidence intervals were calculated using the Wald confidence intervals methods using the function *confint* from the base package *stats*.

#### Survival analysis over ageing

We estimated the change in infection susceptibility either weekly from 7 to 40 days of age, or at 7–10 and 35 days of age. This was done using a Cox proportional hazards model with strata to take into account the fact that experiments were replicated over different days. The model was specified as: coxph(Surv(Time_to_death,Censor) ∼ Age + Age:Sex + strata(Experiment_ID).

#### Comparison of ageing models

To describe the pattern of age-related increase in infection susceptibility, we compared five mathematical models representing different dynamics of biological ageing. 1) The Linear model (i.e., constant ageing) assumes that susceptibility increases at a constant rate with age, following the equation: *HR(x) = α + β × x*; where *HR(x)* is the hazard ratio at age *x*, *α* is the baseline hazard ratio, *β* is the constant rate of increase, and *x* is age. This model represents additive ageing where vulnerability increases by fixed amounts over time. 2) The Gompertz model (i.e., exponential ageing) [104] assumes that mortality risk increases exponentially with age, following the equation: *HR(x) = α * exp(β * x)*; where *HR(x)* is the hazard ratio at age *x*, *α* is the baseline hazard ratio, *β* is the ageing rate parameter, and *x* is age. The model was fitted in its log-linear form, log(HR) = log(α) + βx. Taking the natural logarithm linearises this relationship: *log[HR(x)] = log(α) + β × x*. This model represents a case where damage accumulates multiplicatively over time and is the classical ageing theory for the dynamics of overall mortality risk. 3) The Weibull model (i.e., power-law ageing) [139] assumes that ageing follows a power-law relationship with age, following the equation: *HR(x) = α x^β*; where β is the shape parameter determining the ageing pattern. The model was fitted in its log-linear form, log(HR) = log(α) + β log(x). Taking the natural logarithm linearises this relationship: *log[HR(x)] = log(α) + β × log(x)*. This model captures flexible ageing dynamics commonly observed in biological systems. 4) The Logarithmic model (i.e., ageing with asymptotic plateau) assumes that ageing effects decelerate over time, with the relationship: *HR(x) = α + β × log(x)*. This pattern suggests “early” vulnerability followed by adaptation or selection effects that reduce the rate of ageing progression, or simply that ageing reaches a maximum vulnerability. 5) The Logistic model (i.e., ageing with threshold) represents sigmoidal ageing with distinct phases, with the relationship: *HR(x) = α + (β - α) / [1 + exp(-k(x - x₀))]*; where *α* is the lower asymptote, *β* is the upper asymptote, *k* is the steepness parameter, and *x₀* is the inflection point. This model suggests initial stability, followed by rapid ageing, then a plateau at maximum vulnerability. The Linear, Logarithmic, and Logistic models represent additive ageing processes (i.e., constant increases in hazard ratio), whilst the Gompertz and Weibull models represent multiplicative ageing processes (i.e., proportional increases in hazard ratio).

To fit the five models, hazard ratios were calculated from Cox proportional hazards models for each age group relative to the youngest age group, analysed separately by sex (data from Figure 1A’). The analysis combined data from both sexes and all experimental replicates, with each sex-replicate combination treated as an independent unit through mixed-effects modelling using Sex_Replicate as a random effect (e.g., Female_Rep1, Male_Rep1, etc.). This approach accounted for experimental structure whilst testing universal ageing patterns and assumes that the ageing pattern (i.e., rate and shape of change) is universal across sex-replicate combinations but allows for baseline differences in vulnerability levels. We did not have the statistical power to make the same comparative analysis within each sex. For the Linear model, we fitted mixed-effects models using *lmer()* with hazard ratios against age and random intercepts by Sex_Replicate. For the Gompertz model, we fitted mixed-effects models using *lmer()* with log-transformed hazard ratios against age and random intercepts by Sex_Replicate. For the Weibull model, we fitted mixed-effects models using *lmer()* with log-transformed hazard ratios against log-transformed age and random intercepts by Sex_Replicate. For the Logarithmic model, we fitted mixed-effects models using *lmer()* with hazard ratios against log-transformed age and random intercepts by Sex_Replicate. For the Logistic model, we fitted non-linear mixed-effects models using *nlme()* with SSlogis function and random effects on the asymptote parameter (maximum HR) by Sex_Replicate. All models were checked for proper convergence.

We used AIC as the primary model selection criterion, which balances goodness of fit against model complexity (AIC = 2k - 2ln(L)), where k is the number of parameters and L is the likelihood. The Gompertz and Weibull models are fitted to log-transformed hazard ratios, whereas the Linear, Logarithmic and Logistic models are fitted on the natural scale, so their AIC values are not directly comparable without adjustment. We therefore applied a Jacobian correction (adding 2 Σ log(HR)) to the log-scale models, placing all five on a common scale. We also excluded the hazard ratios at 7 days of age, which are fixed at 1 by construction as the reference category rather than estimated, before fitting. Lower AIC values indicate better models. We calculated ΔAIC = AIC_i - AIC_min for each model considering that ΔAIC < 2 is a substantial support for the model.

We also investigated the goodness of fit by calculating R^2^ values for each model to assess the proportion of variance explained by fixed effects (marginal R^2^). For linear mixed-effects models (Linear, Gompertz, Weibull, Logarithmic), we used the *r.squaredGLMM()* function from the *MuMIn* package. For non-linear mixed-effects models (logistic), we calculated R^2^ as the squared correlation between fitted and observed values.

We then extracted parameters from the best-fitting model for biological interpretation. For the Weibull model: Shape parameter (*β*): The power-law exponent determining ageing dynamics. When *β* < 1, ageing decelerates (i.e., early vulnerability with slower later increases); when *β* > 1, ageing accelerates; when *β* = 1, the hazard ratio increases in direct proportion to age.

#### Infection susceptibility at young and old age in DGRP lines

We estimated the change in infection susceptibility at 10 and 35 days of age in several DGRP lines across four experiments, each containing 7 or 8 lines, with one line (i.e., RAL-821) included in every experiment. RAL-821 was the reference that allowed comparisons across experiments. We used a Cox proportional hazards model to consider this experimental design. The model was specified as: coxph(Surv(Time_to_death,Censor) ∼ Line + Line : Age_at_infection + strata(Experiment_ID)). This was performed for each sex independently, as we were not aiming at testing the triple interaction, but to calculate the hazard ratios related to each age group and their differences (i.e., age-related increased susceptibility).

#### Correlation analyses between sexes and across populations

To estimate the HR characterising the relative lifespan of each of the 12 lines, we used the same approach as for studying their susceptibility. The model was specified as: coxph(Surv(Age,Censor) ∼ Line + strata(Experiment_ID), with RAL-821 as reference. This yields population-specific HRs relative to RAL-821 within each block, controlling for systematic block effects.

Correlations between male and female lifespan were tested using linear regression with block as covariate, where beta values represent the log(HR) relative to RAL-821. This explicitly controls for block variation while testing sex correlations. Robustness was confirmed by repeating the analyses with Block 1 excluded.

Because the model uses RAL-821 as the reference level (i.e., intercept), all other population estimates represent contrasts relative to RAL-821. Including RAL-821 – which by definition has β = 0 for both traits – would artificially inflate correlations due to non-independence and violate the statistical assumption of independence among the populations. Therefore, RAL-821 was excluded from all correlation analyses in which estimates are expressed relative to it. This does not apply to Figures 3F and 4B, where each population is compared with its own young baseline rather than with RAL-821: in these analyses RAL-821 has an estimated, non-zero value like every other population, and is included.

For bivariate correlations between the sexes (Figures 3B, 3D, 3F and 4B), regressions were fitted with the male estimate as the response, matching the orientation of the plotted axes, and Pearson correlation coefficients are reported with their 95 % confidence intervals.

Because the population-level estimates differ substantially in precision, the correlation in Figure 3B was additionally fitted with inverse-variance weighting, the same measurement-error weighting applied to PLUD estimates above, and by robust MM-estimation (*lmrob*, package *robustbase*, KS2014 setting), both using all populations. Leave-one-out refits, Cook’s distance and DFBETAS (thresholds 4/n and 2/√n) and robustness weights are reported in Supplementary Material S1, Section 3.2.2.1.

One property is reported descriptively for each population and is not used to exclude data: the divergence between the sexes, defined as the absolute difference between the male and female estimates. In Figure 3B, RAL-443 and RAL-907 are highlighted as the populations with by far the largest divergence between the sexes; in Figure 4B, RAL-517 and RAL-109 are highlighted because the sexes change in opposite directions with age. All are included in the corresponding correlations and regressions.

Populations were omitted from an analysis entirely only on prospective, precision-based criteria (i.e. insufficient sample size to estimate the quantity concerned, or absence of data at one age) which are stated for each figure. On these grounds, RAL-129 and RAL-879 were omitted from male PLUD analyses, and RAL-738 males from analyses at 35 days.

#### Convergence of susceptibility between populations with age

To assess how much of the variation in susceptibility present at 10 days persists to 35 days, we regressed the population-specific log(HR) at 35 days on that at 10 days, separately for each sex. Under the null hypothesis that the age-related increase is independent of baseline susceptibility, this slope is expected to equal 1, so we tested the fitted slope against 1 rather than against 0. Because both variables derive from the same populations, we confirmed by simulation that a slope below 1 cannot be produced by measurement error or regression to the mean alone. Populations lacking an estimate for either sex at either age were excluded, since the comparison requires both sexes for the same population. Details are in Supplementary Material S1, Section 3.2.5.

#### PLUD estimates

PLUD was estimated across multiple dilutions and duplicates, limited to cases where individual plate counts were less than 150 colony-forming units (CFUs), using maximum-likelihood estimation with left-truncated Poisson models. Individual estimates were then screened with a Rosner test applied within each Line × Age × Block × Sex group (Line × Age × Replicate × Sex for the age series), following Duneau et al. [42]. In an experiment of this design a small number of flies die of causes unrelated to the infection, and the pathogen load at death of such individuals has no biological meaning; because the proportion of these deaths can differ between the groups being compared, retaining them can generate spurious differences, as Duneau et al. demonstrate by simulation. The screen removed fewer than 5 % of individual estimates in both experiments, distributed across groups and evenly divided between young and old flies (Supplementary Material S1, Sections 4.1 and 4.5). Estimates were also excluded where the maximum-likelihood estimation failed to converge, or where total CFU counts were 10 or fewer, since such a load cannot represent the maximum bacterial burden a host can sustain and is therefore not a PLUD by definition. Both criteria exclude measurements that cannot be interpreted as a PLUD, rather than measurements that are simply extreme.

For the population comparison experiment, these PLUD estimates were modelled to obtain population-specific estimates – relative to RAL-821 – for each sex separately, using a mixed-effect generalised linear model implemented in the *spaMM* package [140]. Since bacterial loads are estimated through serial dilution, some estimates are more precise than others. To account for this heterogeneous measurement uncertainty, observations with less precise estimates (i.e., higher estimate variance) were given less weight in the model fit. The measurement error variance for each observation was calculated as the square of the standard error from the maximum likelihood estimate and incorporated into the model using the *resid.model* argument in the *fitme* function, which specifies known heterogeneous residual variance. ‘Population’ and ‘Age’ were fitted as fixed effects, while ‘Experiment ID’ was fitted as a random effect to account for block effects. The model was specified as: log10(Bacterial load per fly) ∼ Population + Population:Age + (1|Experiment ID), resid.model = ∼ Var_logload, where Var_logload represents the measurement error variance.

For the age-by-sex experiment, the same maximum-likelihood estimation and variance-weighting approach was used. In this experiment, a single genotype was tested with both sexes across ages, allowing PLUD to be estimated across the combination of age and sex. The model was specified as: log10(Bacterial load per fly) ∼ Age * Sex + (1|Experiment ID), resid.model = ∼ Var_logload. Confidence intervals for all estimates were calculated using parametric bootstrap resampling (n = 999 iterations) with the percentile method.

## List of Abbreviations

AIC: Akaike Information Criterion
AMP: Antimicrobial Peptide
AOx: Ashworth Outcrossed
BBSRC: Biotechnology and Biological Sciences Research Council
CFU: Colony Forming Unit
CI: Confidence Interval
ΔAIC: Delta AIC
DGRP: Drosophila Genetic Reference Panel
DTP: Doctoral Training Programme
GLMM: Generalised Linear Mixed Model
HR: Hazard Ratio
HTML: Hypertext Markup Language
IMD: Immune Deficiency Pathway
LB: Luria Bertani
NF-kB: Nuclear Factor kB
NIH: National Institutes of Health
OD600: Optical Density at 600 nm
PBS: Phosphate Buffer Saline
PLUD: Pathogen Load Upon Death
RAL (Prefix): Raleigh
ROS: Reactive Oxygen Species
RRID: Research Resource Identifier
UKRI: UK Research and Innovation

## Funding

This work was supported by a UKRI-BBSRC EastBio DTP studentship to BMB; a Darwin Trust studentship to RLB; a Fundação para a Ciência e a Tecnologia (FCT) research contract (2023.08149.CEECIND), FCT (UID/00329/2025) and FCT CHANGE (LA/P/0121/2020) grants to DFD; and start-up funding from The University of Edinburgh, a Wellcome Trust (210183/Z/18/Z) and a Royal Society grant (RGS\R1\221328) to JCR.

## Acknowledgements

We wish to thank Darren Obbard, Pedro Vale, Neha Agrawal and their lab members for fruitful discussions around this work, Pedro Vale for the AOx line and Katy Monteith and Mariangel González for its maintenance, Nicolas Buchon and Jean-Baptiste Ferdy for discussion of data and advice on infections, and Jarrod Hadfield and Jean-Baptiste Ferdy for advice on statistical analyses. We wish to thank the three anonymous reviewers who helped to greatly improve the final version of the manuscript.

---

**Supplementary Figure S1.**
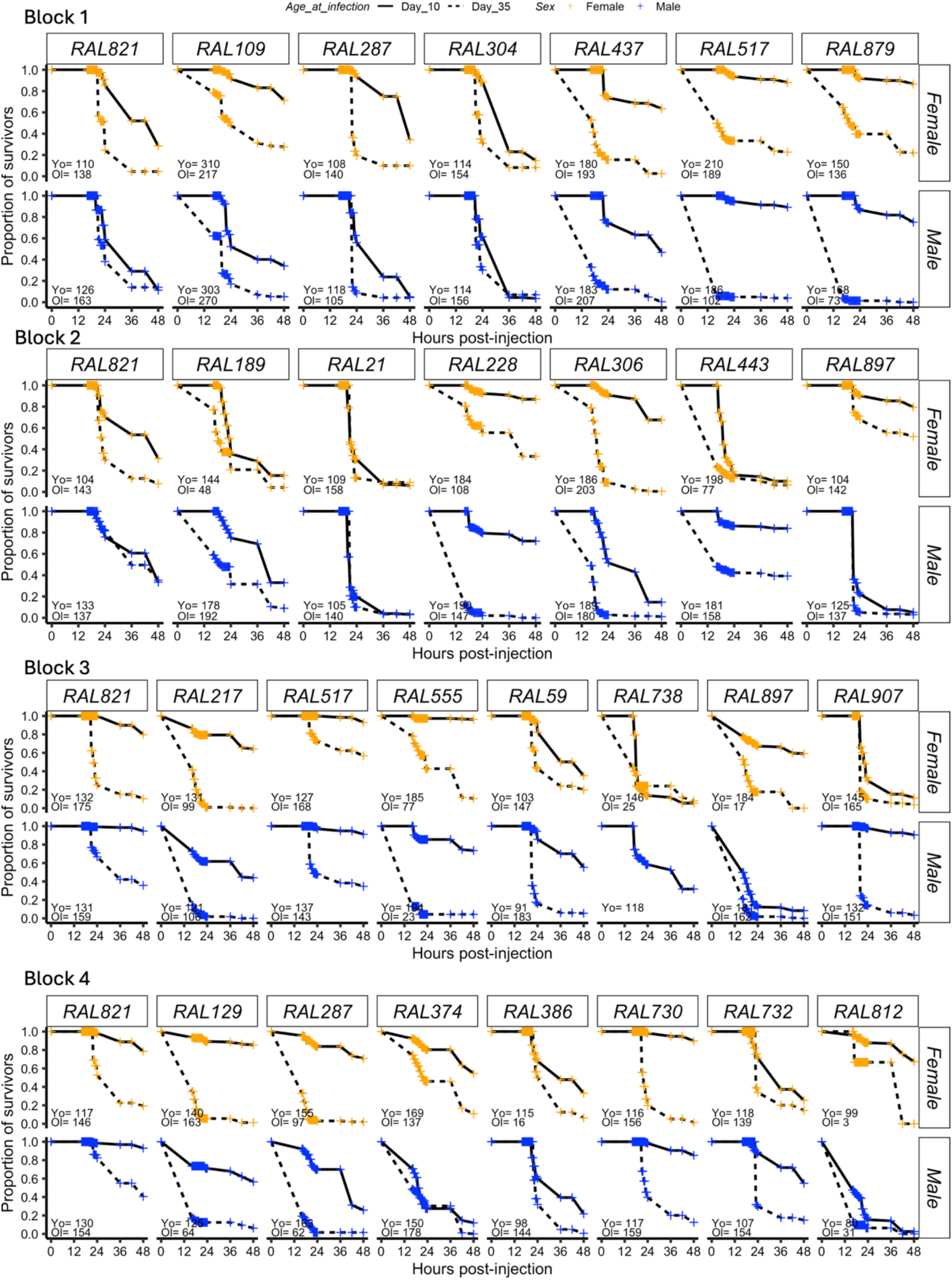
Survival of young (10 days) and old (35 days) males and females from 22 DGRP lines.

**Supplementary Figure S2.**
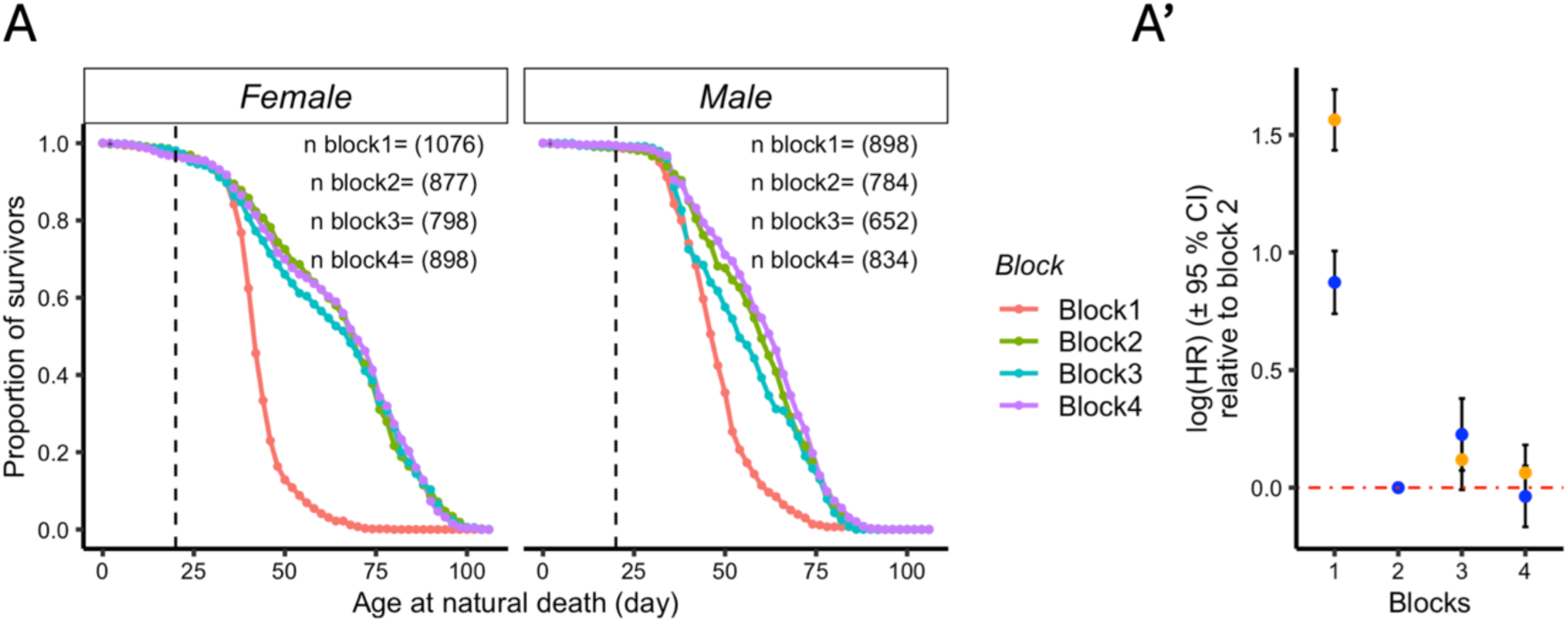
Repeatability of lifespan across blocks. Survival curves and log(Hazard ratio) of RAL821 in each block. Lifespan was shorter in block 1, but the three other blocks were equivalent.

**Supplementary Figure S3.**
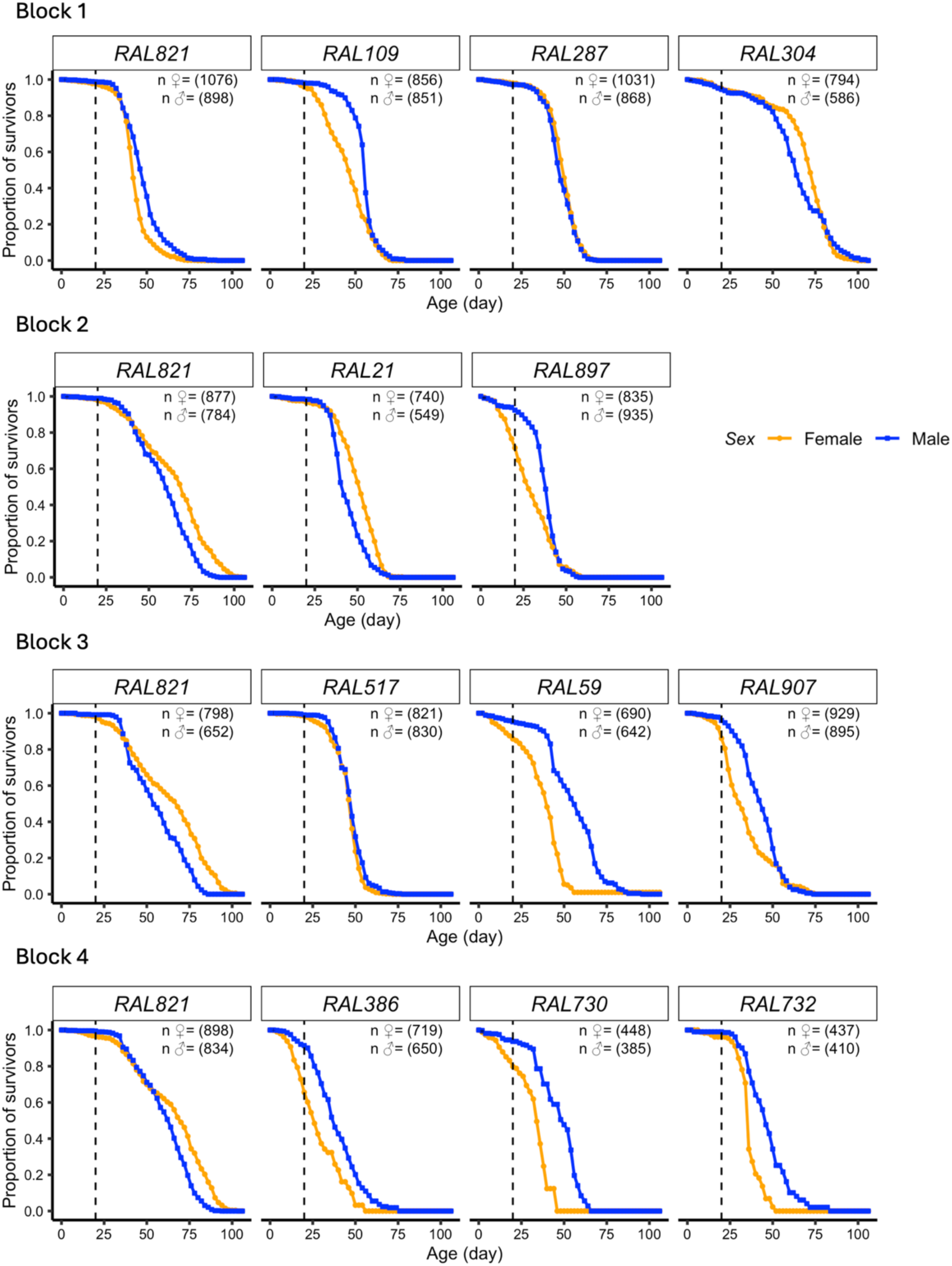
Survival curves of both sexes of 12 DGRP lines across blocks.

**Supplementary Figure S4.**
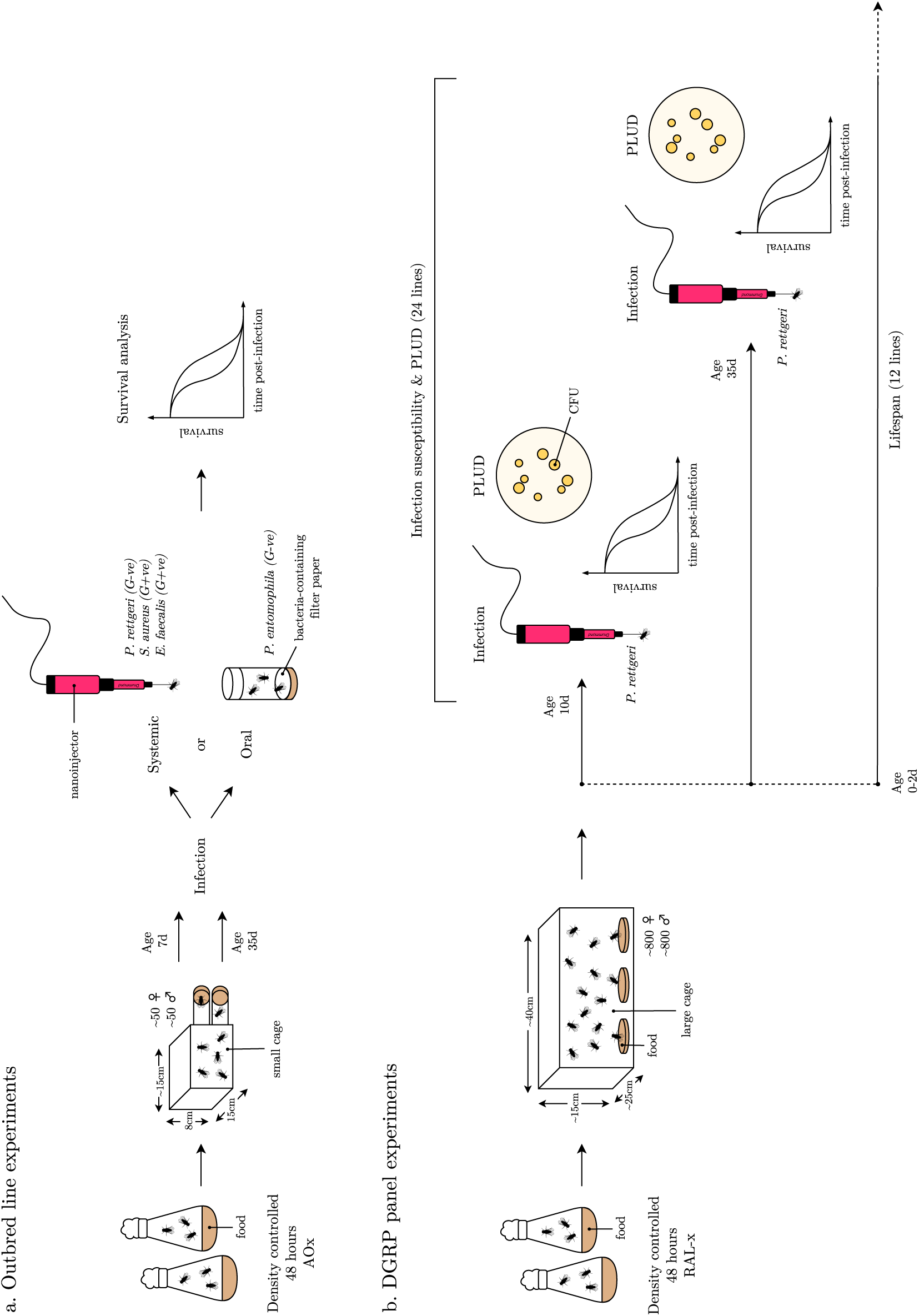
Schematic representation of experimental design, including *D. melanogaster* populations and pathogen strains.

**Supplementary Table S1.** *Drosophila* Genetic Reference Panel (DGRP) lines used in this study.

| DGRP Line | Synonym | RRID |
| --- | --- | --- |
| DGRP-21 | RAL-21 | RRID:BDSC_28122 |
| DGRP-59 | RAL-59 | RRID:BDSC_28129 |
| DGRP-109 | RAL-109 | RRID:BDSC_28140 |
| DGRP-129 | RAL-129 | RRID:BDSC_28141 |
| DGRP-189 | RAL-189 | RRID:BDSC_28152 |
| DGRP-217 | RAL-217 | RRID:BDSC_28154 |
| DGRP-228 | RAL-228 | RRID:BDSC_28157 |
| DGRP-287 | RAL-287 | RRID:BDSC_28165 |
| DGRP-304 | RAL-304 | RRID:BDSC_25177 |
| DGRP-306 | RAL-306 | RRID:BDSC_37525 |
| DGRP-374 | RAL-374 | RRID:BDSC_28185 |
| DGRP-386 | RAL-386 | RRID:BDSC_28192 |
| DGRP-437 | RAL-437 | RRID:BDSC_25194 |
| DGRP-443 | RAL-443 | RRID:BDSC_28199 |
| DGRP-517 | RAL-517 | RRID:BDSC_25197 |
| DGRP-555 | RAL-555 | RRID:BDSC_25198 |
| DGRP-730 | RAL-730 | RRID:BDSC_25202 |
| DGRP-732 | RAL-732 | RRID:BDSC_25203 |
| DGRP-738 | RAL-738 | RRID:BDSC_28223 |
| DGRP-812 | RAL-812 | RRID:BDSC_28240 |
| DGRP-821 | RAL-821 | RRID:BDSC_28243 |
| DGRP-879 | RAL-879 | RRID:BDSC_28254 |
| DGRP-897 | RAL-897 | RRID:BDSC_28260 |
| DGRP-907 | RAL-907 | RRID:BDSC_28262 |

This table lists all DGRP stocks used in our experiments, all of which were obtained from the Bloomington *Drosophila* Stock Center. For each line, the corresponding synonym and
Bloomington Research Resource Identifier (RRID) are provided. No additional stocks were generated for this work.

**Supplementary Table S2.** Pathogenic bacterial strains used in this study.

| Species | Strain | Abbreviation | Source |
| --- | --- | --- | --- |
| *Providencia rettgeri* | Dmel | *P. rettgeri* | Brian P. Lazzaro |
| *Staphylococcus aureus* | PIG1 | *S. aureus* | Brian P. Lazzaro |
| *Enterococcus faecalis* | OG1RF | *E. faecalis* | Stéphane Mesnage |
| *Pseudomonas entomophila* | Dmel | *P. entomophila* | Brian P. Lazzaro |

This table lists all pathogenic bacterial strains used in our experiments. Strains were obtained as gifts from principal investigators in research laboratories rather than from commercial culture collections. For each pathogen, the species name, strain designation, abbreviation used in the main text, and the providing source are indicated.

## Notes

### Competing Interest Statement

The authors have declared no competing interest.

### Summary of Updates

New modelling and new experimental data have been added to further clarify infection susceptibility over age and the ageing patters therein. In addition, there are changes to the text, figure legends and methods, amendments to statistical analyses of data, and additional supplementary data. Author affiliations have been updated to include ORCiD for all authors where possible.

